## Supplementary figures and images for "Laminin-defined Mechanical Status Modulates Retinal Pigment Epithelium Functionality"

### Sup. fig. 1

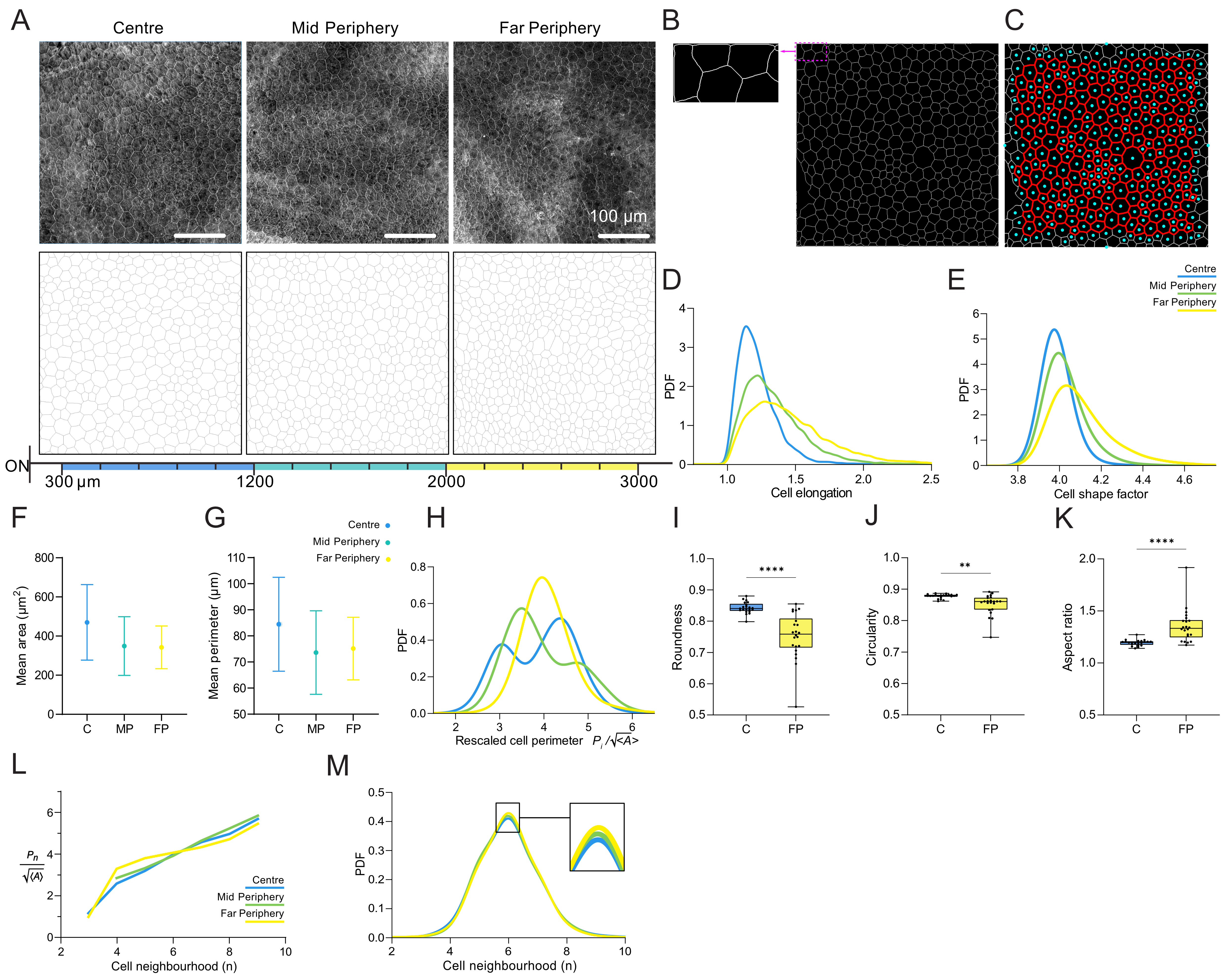

### Sup. fig. 2

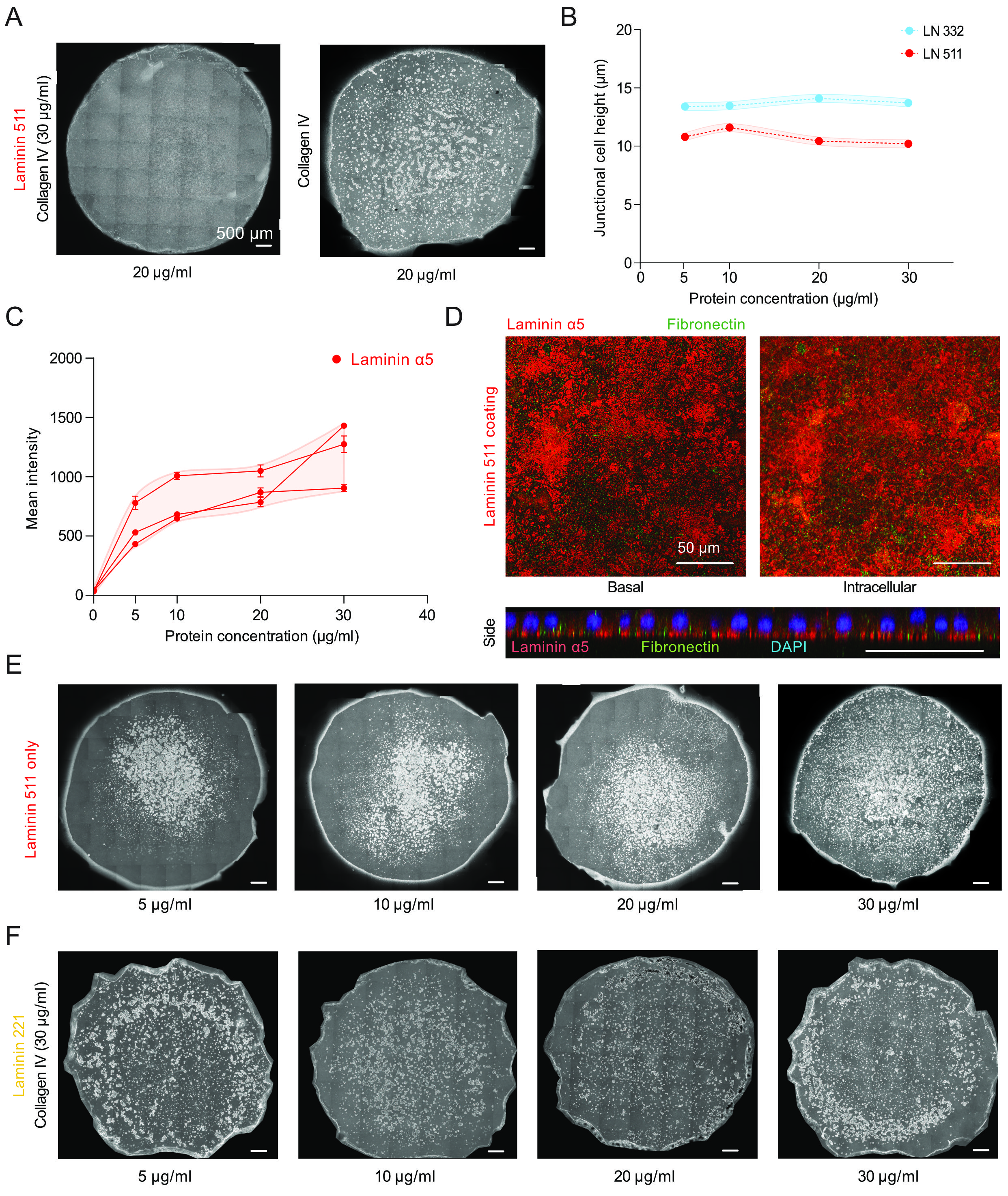

### Sup. fig. 3

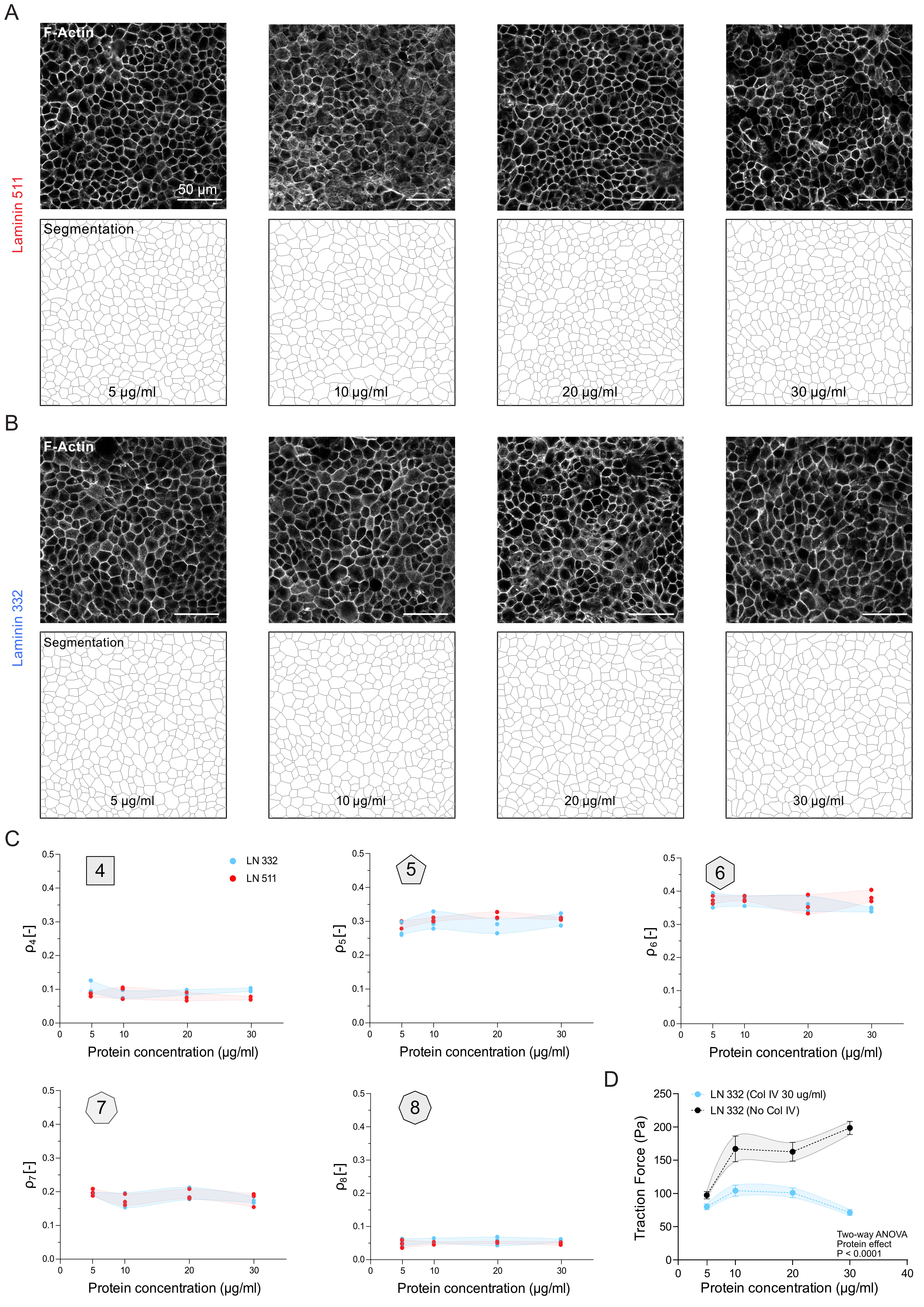

### Sup. fig. 4

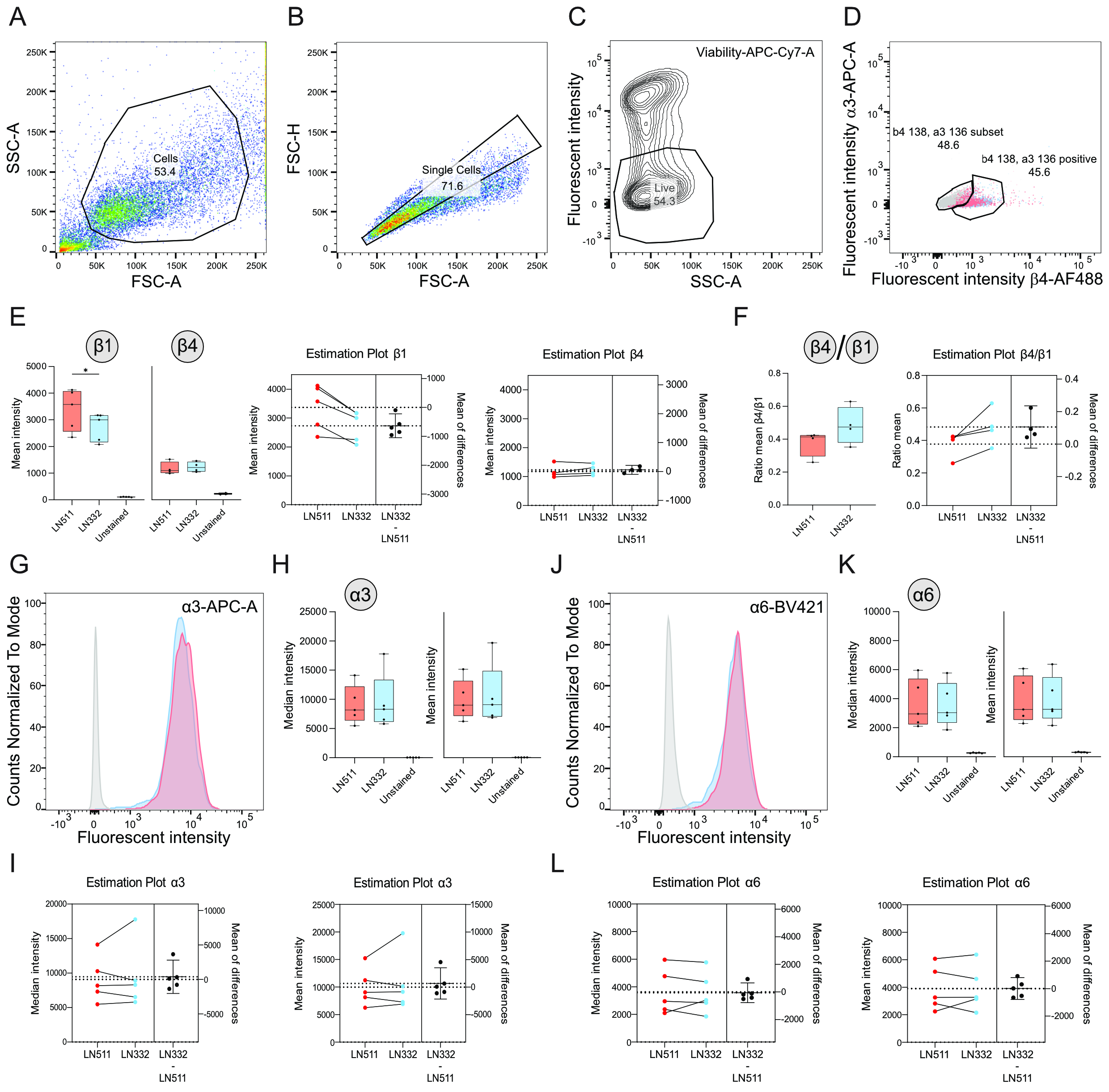

### Table S 2

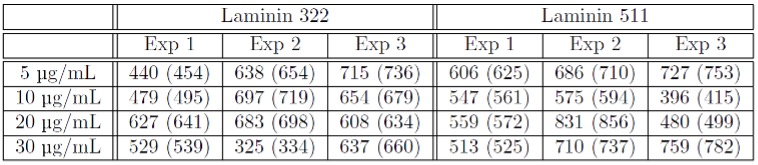
