## Supplementary material for "Laminin-defined Mechanical Status Modulates Retinal Pigment Epithelium Functionality": Table S 1

| Name | Origin | Source | Concentration (µg/ml) / Dilution |
|---|---|---|---|
| **Primary antibodies** | | | |
| Anti mouse laminin 111 (pan-Laminin) | rabbit | Gift from Lydia Sorokin | Rabbit serum, 1:2000 |
| Anti mouse laminin α1 (clone 200) | rat | Gift from Lydia Sorokin | Hybridoma supernatant |
| Anti mouse laminin α2 (clone 4H8-2) | rat | Gift from Lydia Sorokin | Hybridoma supernatant |
| Anti human laminin 332 | rabbit | Gift from Monique Aumailley | Rabbit serum, 1:2000 |
| Anti mouse laminin α4 | rabbit | Gift from Lydia Sorokin | Rabbit serum, 1:1000 |
| Anti mouse laminin α5 | rabbit | Gift from Lydia Sorokin | Rabbit serum, 1:1000 |
| Anti mouse collagen type IV | mouse | Merck Millipore, AB756P | 12.5 |
| Anti mouse collagen type I | rabbit | Novusbio, NB600-408 | 5 |
| Anti mouse elastin | rabbit | Abcam, ab21610 | 5 |
| Phalloidin iFluor 647 | | Abcam, ab176753 | 1x |
| Phalloidin iFluor 488 | | Abcam, ab176759 | 1x |
| Ant human ZO-1 | rabbit | Thermo Fisher, 61-7300 | 5 |
| Anti human ezrin (clone 3C12) | mouse | Abcam, ab4069 | 20 |
| Anti human fibronectin | rabbit | Sigma, F3648 | 2.5 |
| **Antibodies used for flow cytometry** | | | |
| Integrin β1-PE (clone 12G10) | mouse | Santa Cruz, sc-59827 PE | 1 mg per million cells |
| Integrin β4-Alexa Flour 488 (clone A9) | mouse | Santa Cruz, sc-13543 AF488 | 1 mg per million cells |
| Integrin α3-Alexa Flour 647 (clone P1B5) | mouse | Santa Cruz, sc-13545 AF647 | 1 mg per million cells |
| Integrin α6-Brilliant Violet 421 (clone GOH3) | rat | BioLegend, 313623 | 5 ml per million cells |
| **Secondary antibodies** | | | |
| Anti-mouse-IgG AF488 | goat | Invitrogen, A11001 | 4 |
| Anti-mouse-IgG AF594 | goat | Invitrogen, A11005 | 4 |
| Anti-mouse-IgG AF647 | goat | Jackson/Dianova, 111-605-144 | 10 |
| Anti-rabbit-IgG AF594 | goat | Invitrogen, A11012 | 4 |
| Anti-rabbit-IgG AF647 | goat | Molecular Probes, A21235 | 10 |
| Anti-rat-IgG AF555 | goat | Invitrogen, A21434 | 4 |
| Anti-rat-IgG AF647 | goat | Thermo Fisher, A21247 | 10 |
